# A general Bioluminescence Resonance Energy Transfer (BRET) protocol to measure and analyze protein interactions in mammalian cells

**DOI:** 10.1101/2024.07.05.602189

**Authors:** Carla Jane Duval, Candy Laura Steffen, Karolina Pavic, Daniel Kwaku Abankwa

## Abstract

Bioluminescence resonance energy transfer (BRET) allows to quantitate protein interactions in intact cells. Here we provide a step-by-step protocol for measuring BRET due to transient interactions of oncogenic K-RasG12V in plasma membrane nanoclusters of HEK293-EBNA cells. We describe how to seed, transfect and replate cells, followed by their preparation for BRET-measurements on a microplate reader and detailed data analysis steps. For details on how to apply this protocol, please refer to Steffen et al., 2024 ^1^.

**Graphical abstract:** 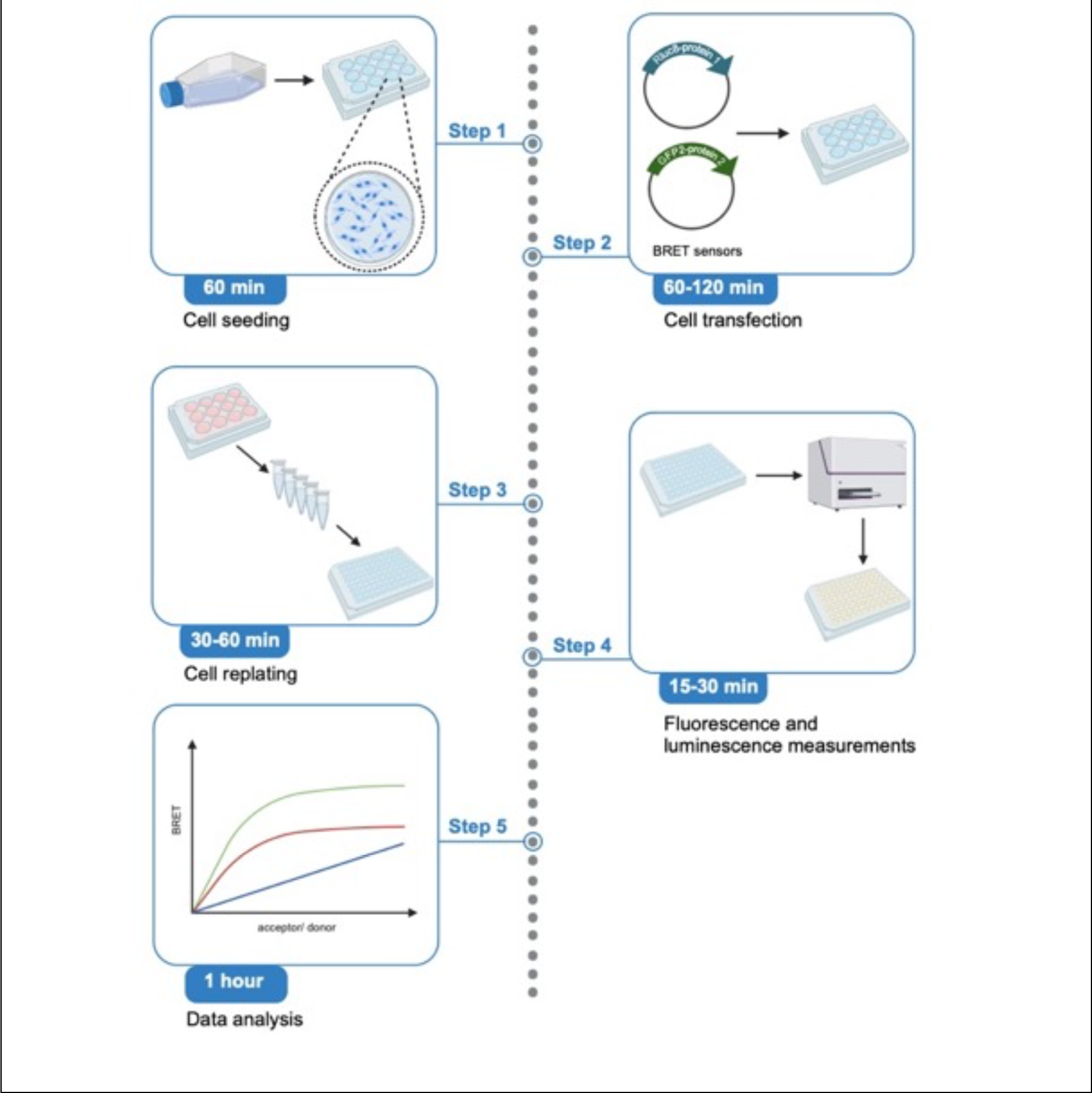

## Before you begin

### Background

Bioluminescence resonance energy transfer (BRET) enables quantitative molecular interaction experiments in the native cellular environment. BRET has thus been used to investigate diverse protein interactions in the cytosol, nucleus and cellular membranes ^2^.

BRET refers to the radiation-free transfer of the donor energy, provided in our case by the Renilla luciferase mutant RLuc8-catalyzed conversion of the luciferase substrate coelenterazine 400a, to the acceptor, green fluorescent protein 2 (GFP2). The donor and acceptor together form the BRET-pair, for which various combinations have been established, including more recently NanoLuc and mNeonGreen^3^. For BRET to occur several conditions must be fulfilled, such as an overlap of the emission spectrum of the donor and excitation spectrum of the acceptor ^2^. Furthermore, donor and acceptor must be in molecular proximity (here 4 - 12 nm) for a change in the BRET-signal, which is the emission by the acceptor after donor excitation ^4^.

By genetically fusing proteins or their domains to donor and acceptor, BRET-biosensors can be constructed, where the interaction is mediated by the fused proteins. As an example, we use the Rluc8- K-RasG12V/ GFP2-K-RasG12V BRET-biosensor, where BRET emerges due to the transient approximation of K-RasG12V in proteo-lipid complexes at the plasma membrane, called nanocluster ^5^. Design and optimization of such biosensors can be challenging and was discussed elsewhere ^6^.

Our protocol provides detailed instructions on conducting donor saturation-titration BRET experiments on a conventional microplate reader that can detect both fluorescence and luminescence. In these experiments the BRET-ratio is plotted as a function of the acceptor/ donor-ratio, where a constant amount of RLuc8-tagged donor-construct is co-expressed with increasing amounts of GFP2-tagged acceptor-construct. Classically, the BRETmax- and BRET50-values are determined from fitting a saturation function to the data. Both values essentially characterize the strength or probability of the interaction ^7^. However, true saturation is typically not reached in cells and we therefore introduced the BRETtop value, which is the highest BRET-ratio within a defined range of acceptor/ donor-expression signal ratios ^8^.

### Preparation

1. To get started, prepare mammalian expression vectors encoding donor- and -acceptor constructs. New constructs require significant optimization and testing for uncompromised biological activity. Our example BRET-biosensor, RLuc8-K-RasG12V/ GFP2-K-RasG12V, and its construction by multi-site gateway cloning was previously described by us ^9^.
2. Prepare cell culture medium and coelenterazine 400a substrate needed for the assay.
3. Verify that your fluorescence and luminescence microplate reader has an injector for dispensing microliter amounts of luciferase substrate to 96-well plates. Furthermore, the instrument needs to allow the definition of three measurement channels, for the donor-, acceptor- and BRET-signal. We here describe the setup and operation using a CLARIOstar Microplate Reader for the RLuc8/ GFP2 BRET-pair.

## Key resources table

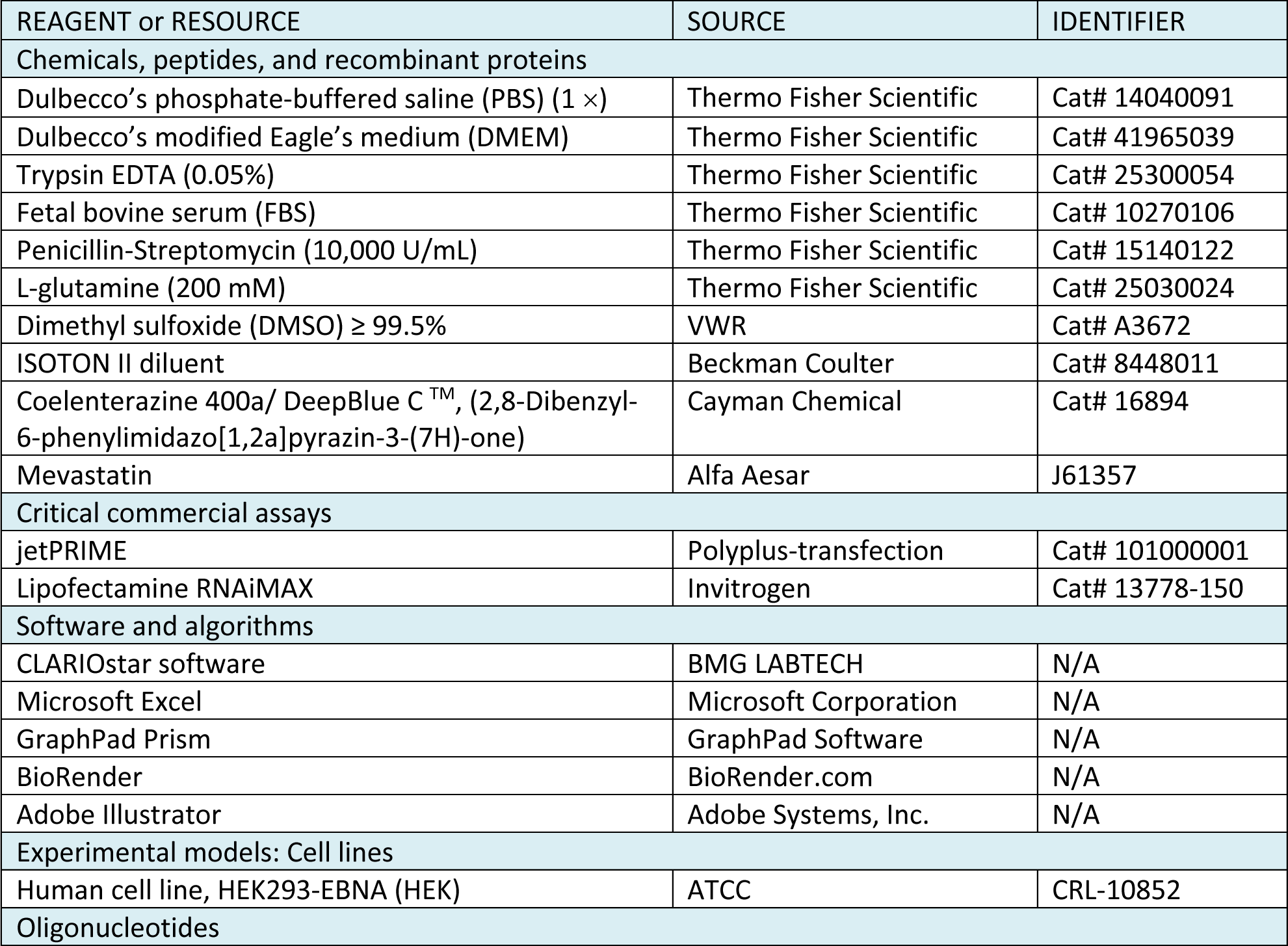
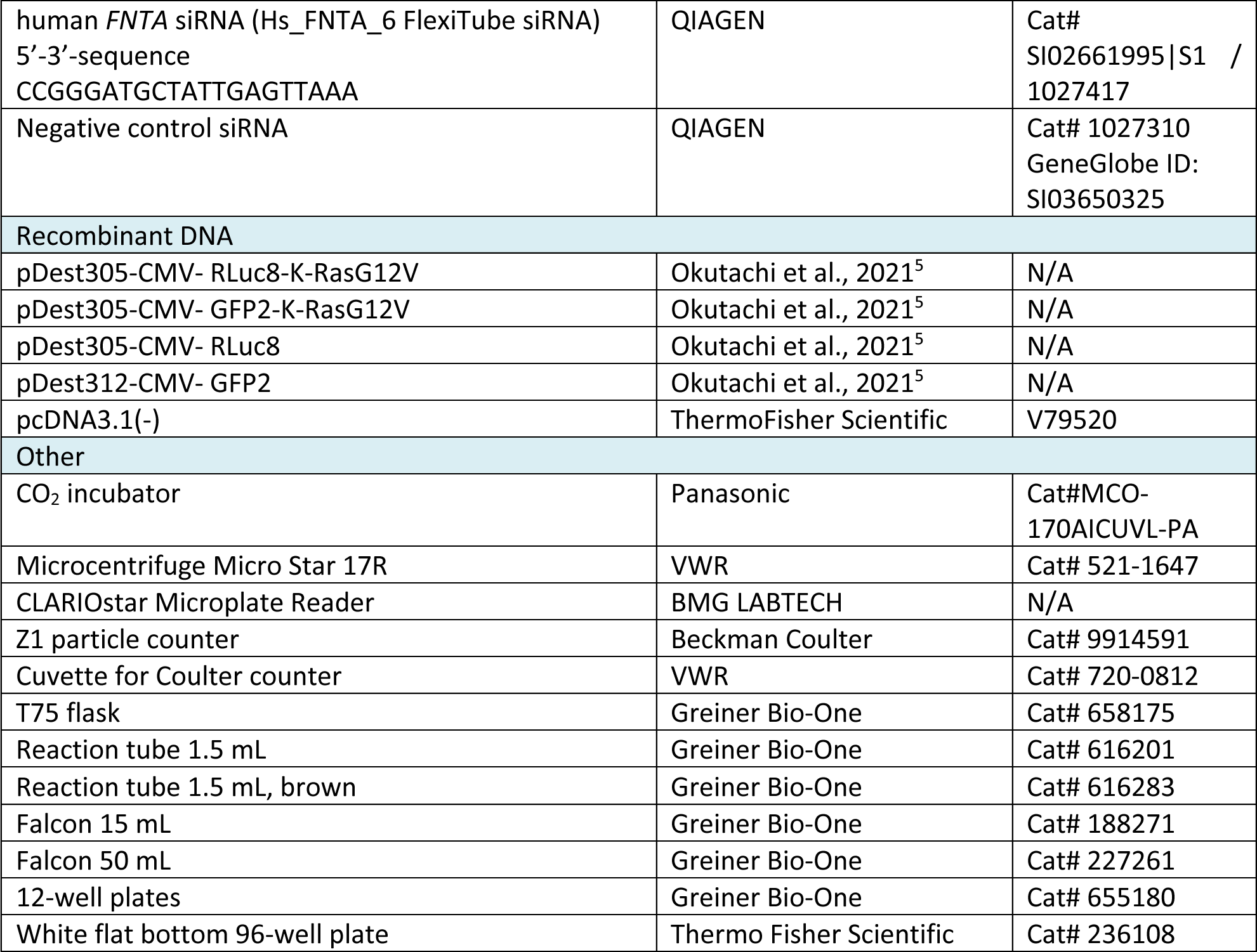

### Materials and equipment setup

- Prepare the growth medium, which is DMEM supplemented with 10% FBS, 2 mM L-glutamine, and 100 U/ mL penicillin/ streptomycin. Store at 4°C for up to 1 month.

**Note:** The addition of penicillin/ streptomycin is recommended to avoid cell culture contamination.

- Prepare a stock solution of the luciferase substrate coelenterazine 400a in 100% ethanol to a final concentration of 1 mM. The stock can be stored in brown reaction tubes at -30°C for several months.

**Note:** Coelenterazine 400a is sensitive to oxidation, therefore, dissolve it fresh before use and keep protected from light.

## Step-by-step method details

### Part 1: Cell seeding

**Timing:** 60 min

This part describes the preparation of a HEK293-EBNA cell culture for the transfection.

1. Prewarm DMEM, PBS and trypsin EDTA in a 37°C water bath.
2. Prepare the cell counter, here Beckman Coulter Z1 Counter, by flushing it twice with Milli-Q water, followed by two flushes with ISOTON II Diluent before use.
3. Grow HEK293-EBNA cells in a T75 flask under humidified 5% CO2 at 37 °C in a complete growth medium until they reach 80 - 90% confluency.
4. Aspirate the growth medium and gently rinse the cells once with 5 mL sterile PBS.
5. Aspirate the PBS and detach the cells by adding 4 mL of trypsin EDTA. Incubate at 37°C until the cells have detached (approximately 3 - 5 min).
6. To neutralize the trypsin EDTA, add 8 mL of growth medium and resuspend by pipetting until all the cells have been washed off from the T75 flask bottom.
7. Transfer the cell suspension to a 15 mL Falcon tube and pellet the cells by centrifugation at 200 Ξ g for 3 min at 22°C - 25°C.
8. Aspirate the supernatant and resuspend the cell pellet in 1 mL of fresh growth medium.
9. To measure the cell concentration, dilute 50 µL of the cell suspension in 10 mL ISOTON II Diluent. This will make a 1:200 cell dilution. Before measuring the cell concentration on the cell counter, make sure that the value specifying the cell dilution on the counter is set to 1:200. The number displayed at the end of the measurement will show the number of cells/ mL.
10. Seed 200,000 cells in 1 mL per well of a 12-well plate.
11. Culture the cells until the desired cell confluency for transfection is reached.

**Optional:** To study the impact of a specific gene knock-down on the BRET-biosensor, HEK293-EBNA can be transfected with siRNA after seeding. As an example, consider the knock-down of the alpha-subunit of farnesyl- and geranylgeranyl-transferase I (*FNTA*), as described in **Table S1**. Growth medium containing siRNA and RNA-transfection reagent needs to be removed before transfecting plasmids encoding the BRET-constructs. Therefore, rinse cells gently once by adding 1 mL of PBS warmed to 37°C.

### Part 2: Cell transfection

**Timing:** 60 - 120 min

BRET-biosensor constructs are transfected into the HEK293-EBNA plated in a 12-well plate, so that each well contains one distinct biological sample (e.g. construct-ratio, drug treatment or constructs with mutation of interest). The amounts of example BRET-biosensor constructs that need to be transfected for a donor saturation-titration BRET experiment example are given in **Table 1**.

12. Make sure that the cells are approximately 50 - 60% confluent before the transfection with jetPRIME.

**Table 1.** Example amounts of transfected BRET-biosensor and donor-only (BRET-control) plasmids for donor saturation-titration BRET experiments. “A” refers to the GFP2-tagged acceptor construct (GFP2-K-RasG12V) and “D” to the RLuc8-tagged donor construct (RLuc8-K-RasG12V). The pcDNA3.1- plasmid is used as empty vector to top up the transfected DNA amount to the same total per well.

| well number | A/D ratio | | plasmid amounts/ ng | | | volume/ μL from 100 ng/ μL plasmid stock | | |
| --- | --- | --- | --- | --- | --- | --- | --- | --- |
|  | A | D | A | D | empty vector | A | D | empty vector |
| 1 | 1 | 1 | 25 | 25 | 975 | 0.25 | 0.25 | 9.75 |
| 2 | 4 | 1 | 100 | 25 | 900 | 1 | 0.25 | 9 |
| 3 | 8 | 1 | 200 | 25 | 800 | 2 | 0.25 | 8 |
| 4 | 12 | 1 | 300 | 25 | 700 | 3 | 0.25 | 7 |
| 5 | 16 | 1 | 400 | 25 | 600 | 4 | 0.25 | 6 |
| 6 | 24 | 1 | 600 | 25 | 400 | 6 | 0.25 | 4 |
| 7 | 32 | 1 | 800 | 25 | 200 | 8 | 0.25 | 2 |
| 8 | 40 | 1 | 1000 | 25 | 0 | 10 | 0.25 | 0 |
| 9 BRET-control | 0 | 4 | 0 | 100 | 900 | 0 | 1 | 9 |

**Note:** Other transfection reagents can be used for which specific optimal transfection conditions may apply.

13. Dilute your plasmid stocks to 100 ng/ µL.
14. The total amount of DNA to be transfected in each well is 1025 ng.
15. For each well, prepare one 1.5 mL reaction tube with the DNA mix . As a BRET-control, transfect cells with only the donor construct (**Table 1**).
16. Dilute the appropriate volume of DNA, indicated in the “volume/ µL from 100 ng/ µL stock” column, in 100 µL jetPRIME buffer and mix by vortexing for 10 s.
17. Before using the jetPRIME reagent, mix it by vortexing 1 - 2 s and briefly spin down to collect the droplets possibly retained inside the tube lid.
18. Add jetPRIME reagent at a 1:3 ratio per µg of DNA. For 1025 ng DNA, use 3 µL jetPRIME reagent. **Note:** If using another transfection reagent consult its instruction manual for specific requirements.
19. Incubate at 22°C - 25°C without shaking for 10-15 min before adding the DNA mix dropwise to the corresponding well.
20. Incubate the well-plate with transfected cells in the incubator (37°C, 5% CO2) for up to 48 h of BRET-biosensor expression.

**Optional:** A drug treatment that modulates the BRET-biosensor interaction can be applied to the cells 24 h after transfection for a maximum of another 24 h within this protocol. Here, we treat some BRET-biosensor samples with 5 µM mevastatin in vehicle (0.1% DMSO/ growth medium). Mevastatin is a competitive inhibitor of 3-hydroxy-3-methylglutaryl-Coenzyme A (HMG-CoA) reductase, which catalyzes the rate-limiting step in the synthesis not only of cholesterol but also of prenyl- pyrophosphates. Thus, Ras prenylation is blocked and consequently its plasma membrane anchorage, which also results in the inhibition of the transient interactions associated with nanoclustering. For our example BRET-biosensor, RLuc8-K-RasG12V/ GFP2-K-RasG12V, the mevastatin treatment would therefore reduce the BRET-ratio.

### Part 3: Cell replating to prepare for the measurement

**Timing:** 30 - 60 min (depending on the number of 12-well plates)

Next, each 12-well sample is replated in quadruplicate into a white flat bottom 96-well plate for the measurements on the plate reader.

21. Carefully aspirate the medium from the cells.

**Note:** Handle the plate gently to avoid cell detachment. If extensive cell detachment is observed under a cell culture microscope, we recommend centrifugation of the plate before aspirating the growth medium. Alternatively, detach all cells from the bottom of the 12-well-plate by pipetting and then centrifuge them for 10 min at 900 × g and 4°C in 1.5 mL reaction tubes before directly proceeding to **step 24** followed by a PBS washing step as described in **steps 22,23**.

22. Rinse the cells by adding 1 mL PBS per well, detach the cells by pipetting and collect in 1.5 mL reaction tubes.
23. Pellet the cells by centrifugation for 10 min at 900 × g and 4°C.
24. Aspirate the supernatant.
25. Resuspend the cells in a slight excess of 380 µL of PBS.
26. For each sample, dispense 4 × 90 µL of cell suspension into four adjacent wells of a white flat bottom 96-well plate as quadruplicate technical repeats.

### Part 4: Fluorescence and luminescence measurements on a plate reader

**Timing:** 15 - 30 min (depending on the number of white flat bottom 96-well plates)

In this part, we first describe the setup of the three BRET-pair specific detection channels for the acceptor- (excitation at 405 ± 10 nm, emission at 515 ± 10 nm), donor- (emission at 410 ± 40 nm), and the BRET-signal (emission at 515 ± 40 nm) on the CLARIOstar microplate reader (**Figure 1**). We then explain how to conduct the measurements starting with the acceptor channel, followed by injection of the luciferase substrate and simultaneous acquisition of the donor- and BRET-channel signals.

27. First, set up the three detection channels as “**Test Protocols**” within the CLARIOstar software.

a. Open the CLARIOstar software on the computer that controls the plate reader.
b. Click on “**Microplate**” and “**Manage Protocols**”, which opens the “**Test Protocols**” window.
c. Click on “**New**”, which opens the “**Measurement Method and Mode**” window. Make sure that “**Fluorescence Intensity**” is selected in the “**Measurement Method**” selection and “**Endpoint**” in the “**Reading mode**” selection (feature 1 in **Fixed_Image_1**) and then click “**OK**”. This will open the “**Fluorescence Intensity – Endpoint**” window.
d. In the “**Fluorescence Intensity – Endpoint**” window, open on its **“Basic Parameters”** tab, type in the name of the new protocol “GFP2_Fluorescence” and select for “**Microplate**” “**NUNC 96**” in the drop-down menu, which is the appropriate setting for our white microplates. Adjust this if another plate type is used (feature 2 in **Fixed_Image_1**).
e. In the “**Presets**” tab enter the following monochromator-filter settings for the acceptor-channel (feature 3 in **Fixed_Image_1**):

- Excitation: 405-20
- Dichroic: 462.5
- Emission: 515-20

**Figure 1:**
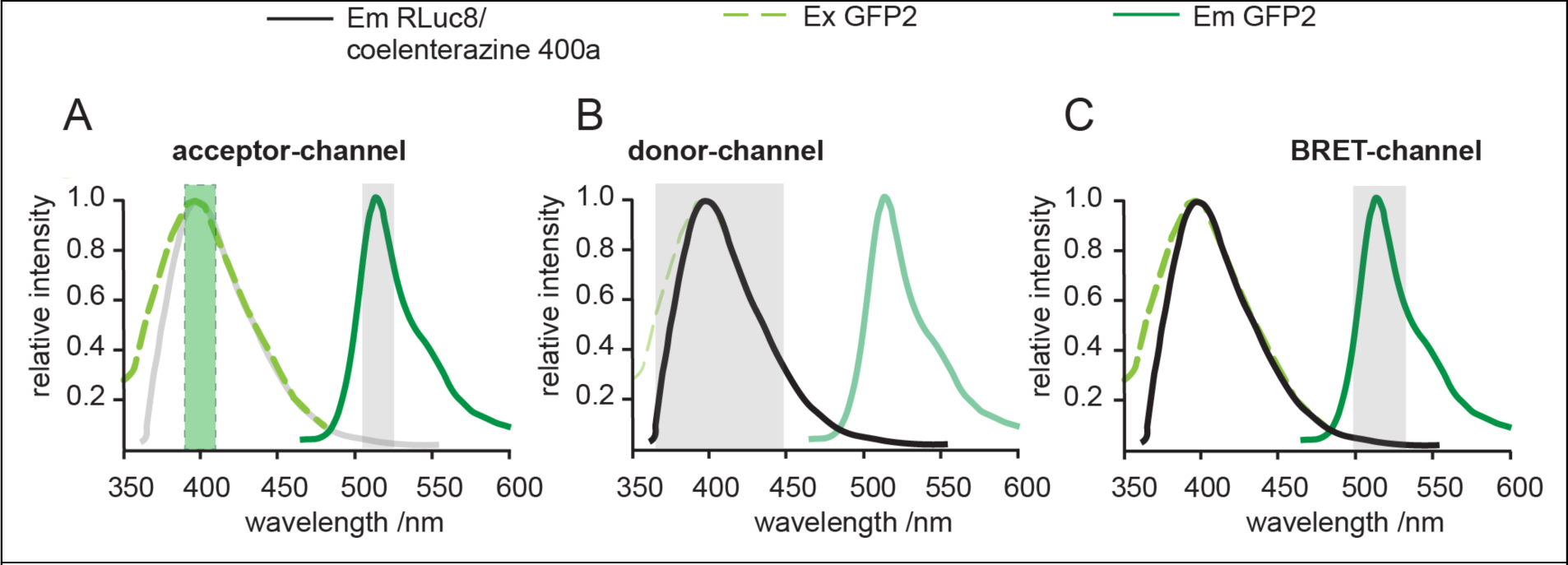
Definition of the three detection channels for BRET-experiments. (**A-C**) The acceptor- (A), donor- (B) and BRET-channels (C) are indicated relative to the emission (Em) and excitation (Ex) spectra of the donor RLuc8/coelenterazine 400a (grey/ black) and the acceptor GFP2 (green). The excitation bandwidth is marked with a light green dashed box (A), while the detection bandwidths are marked with a light grey box (A-C).

and confirm by clicking “**OK**”.

**Note:** The CLARIOstar microplate reader allows to freely chose excitation and emission detection windows, due to its monochromator technology. In the specifications e.g. **“**405-20,” 20 refers to the bandwidth of the monochromator-filter centered at 405 nm, i.e. for detection between 395-415 nm or 405 ± 10 nm. Keep all other features, in the protocol set-up steps not highlighted here at default settings. Specifically, keep “**Optic**” at “**Top optic**”. In the “**General Settings**” field, “**Settling time**” is the time after the microplate moves to the next well and before the measurement begins. It is set to 0.2 s. “**No. of flashes per well**” is set at 50 but can be increased up to 200. All the measurements per flash will be averaged and one intensity value will be obtained per well.

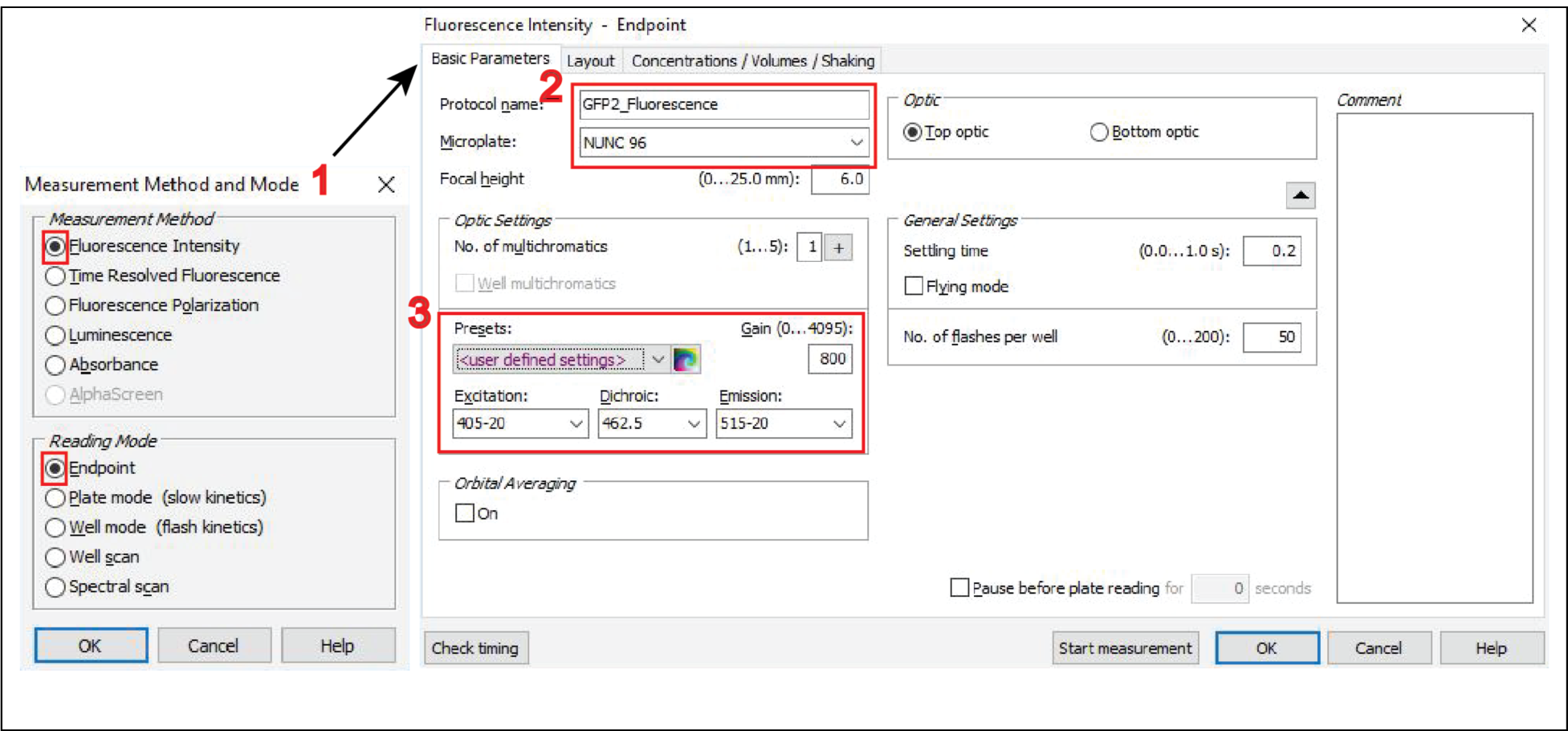

f. Next, create a two-channel protocol for donor- and BRET-channel acquisition, here named “**eBRET2**”. Open another “**Test Protocols**” window and click on “**New**”. Select in the “**Measurement Method and Mode**” window “**Luminescence**” and “**Well mode**” (feature 4 in **Fixed_Image_2**) and then click “**OK**”. By selecting “Well mode”, the luminescence signals will be measured immediately after the injection of the luciferase substrate coelenterazine 400a, with both injection and measurement done well-by- well.
g. In the “**Luminescence – Well mode**” window open on its **“Basic Parameters”** tab, go to the “**Optic Settings**” and change the “**No. of multichromatics**” to “2” (feature 5 in **Fixed_Image_2**).

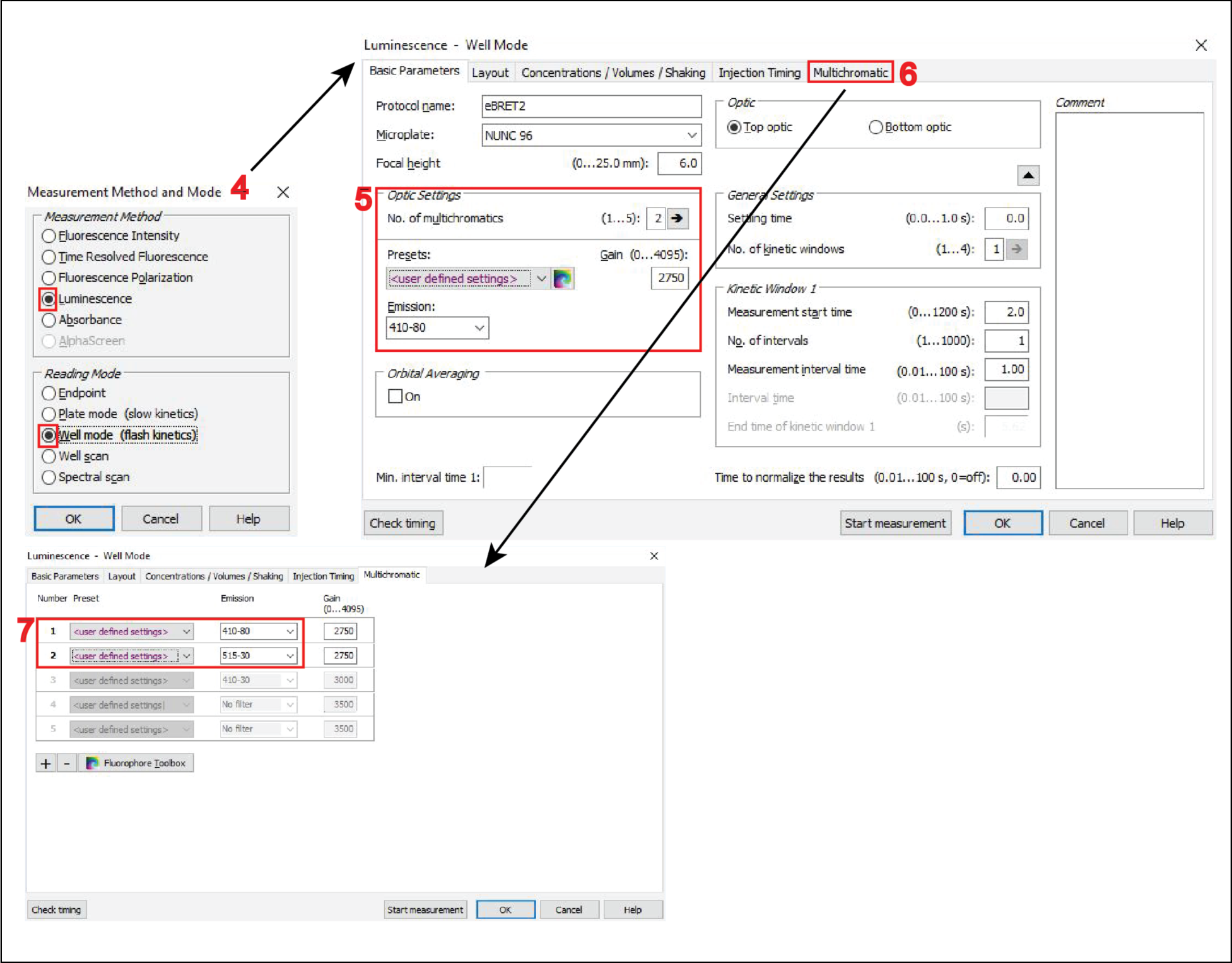

h. In the “**Multichromatic**” tab of the “**Luminescence – Well mode**” window (feature 6 in **Fixed_Image_2**), you then define the donor- and BRET-channel, by entering the two monochromator-filter values “410-80” for the donor- and “515-30” for the BRET- channel (feature 7 in **Fixed_Image_2**).

**Note:** Keep again all other settings, not highlighted here in the protocol set-up steps at displayed default settings. Notably, the “**Settling time**” is set to 0 s. There is no settling time implemented due to the fast substrate conversion.

**CRITICAL:** Before beginning with the actual BRET measurements of the target BRET-biosensor samples, set up optimal gain settings for the photomultiplier tube detector of the microplate reader. See **<u>Troubleshooting 1</u>** for details on how to adjust gain settings on the CLARIOstar Microplate Reader.

28. To start with the measurement of BRET-samples, turn on the CLARIOstar microplate reader and press the button to eject the tray. Place the white flat bottom 96-well plate on it and press the button again to retract the tray.
29. Start the CLARIOstar software from the computer desktop, click on “**Manage Protocols**” and select to display protocols for fluorescence intensity in the “**Test protocols**”.
30. Click on the protocol “GFP2_Fluorescence” to measure the acceptor-channel and click on “**Edit**”. Verify all parameters correspond to those specified in **step 27**.

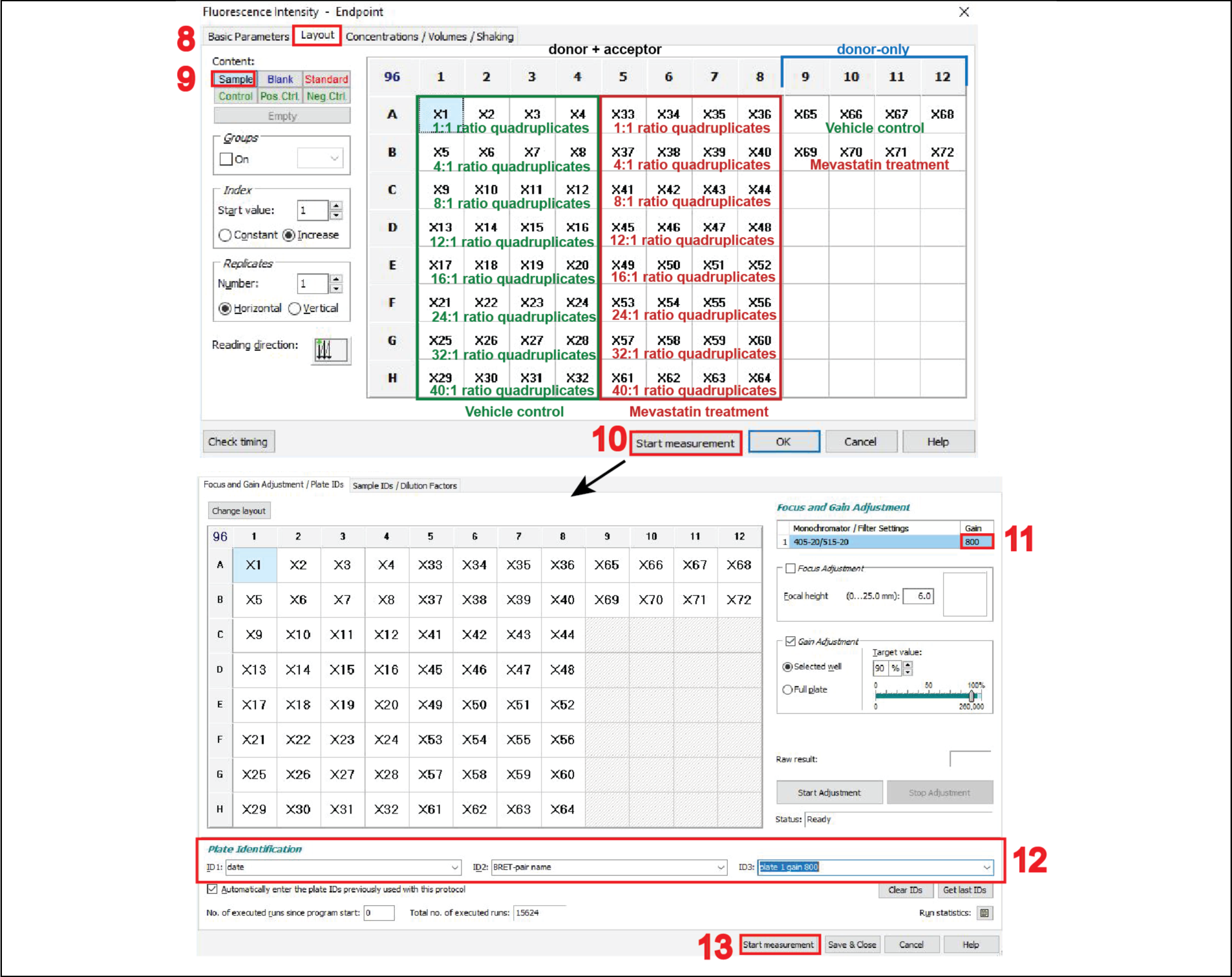

31. This opens the “**Fluorescence Intensity – Endpoint**” window. In the “**Layout**” tab (feature 8 in **Fixed_Image_3**), click on “**Sample**” (feature 9 in **Fixed_Image_3**) and select all wells on the 96- well plate grid that contain samples to be measured.

**Note:** The 96-well plate setup screenshot shows the samples annotated. The vehicle control samples are in green and the mevastatin-treated samples are in red. Plasmid ratios of quadruplicates are also indicated. The donor-only samples are added to the right.

32. The “**Concentrations/ Volumes/ Shaking**” tab is not used in this protocol, therefore keep at default. Click on “**Start measurement**” (feature 10 in **Fixed_Image_3**).
33. In the “**Start measurement”** window, which opens automatically, use the optimal gain settings as determined in **<u>Troubleshooting 1</u>** (feature 11 in **Fixed_Image_3**) and fill out the plate identification ID1, ID2 and ID3 (feature 12 in **Fixed_Image_3**), then click on “**Start measurement**” (feature 13 in **Fixed_Image_3**).

**Note:** Plate identification ID1, ID2 and ID3 are user defined and should be annotated so that the results can be traced back to the corresponding plate and experiment date, e.g. enter ID1: date, ID2: BRET-biosensor, ID3: plate number and gain.

34. For Luminescence measurements, start by preparing a volume of 100 µM coelenterazine 400a appropriate for your sample number by diluting the stock in PBS in a 15 mL Falcon. For a full 96-well plate, prepare 2 mL of coelenterazine 400a.

**Note:** To each well of a white flat bottom 96-well plate, 10 µL of coelenterazine 400a will be added, thus resulting in a final concentration of 10 µM luciferase substrate. Consider that approximately 500 µL of the substrate will be spent when preparing the injection system, described in **step 35**. Additional amounts further account for volume loss when dispensing the substrate from the Falcon into the wells.

35. Prepare the pump and the injection system.

a. Open the lid of the CLARIOstar plate reader, position the input tube end into a Falcon tube with rinsing liquids (below) and place a small beaker underneath the displaced reagent injector for liquid waste collection.
b. Rinse with 100% ethanol by double-clicking the button corresponding to the pump 1, 3 - 4 times (feature 14 in **Fixed_Image_4**).
c. Rinse 3 - 4 times with Milli-Q water
d. Rinse 3 - 4 times with PBS
e. Place the Falcon containing the substrate in the designated place in the instrument and rinse the injection system once with the substrate.
f. Place the reagent injector back in its operating position (feature 15 in **Fixed_image_4**). **Note:** If you have to measure multiple 96-well plates, perform the pump and injection system preparation just before the first injection of coelenterazine 400a. As the substrate precipitates quickly, rinse the injection system with ethanol if the next luminescence reading is more than 20 min later. Consider preparing fresh substrate when precipitates are visible.

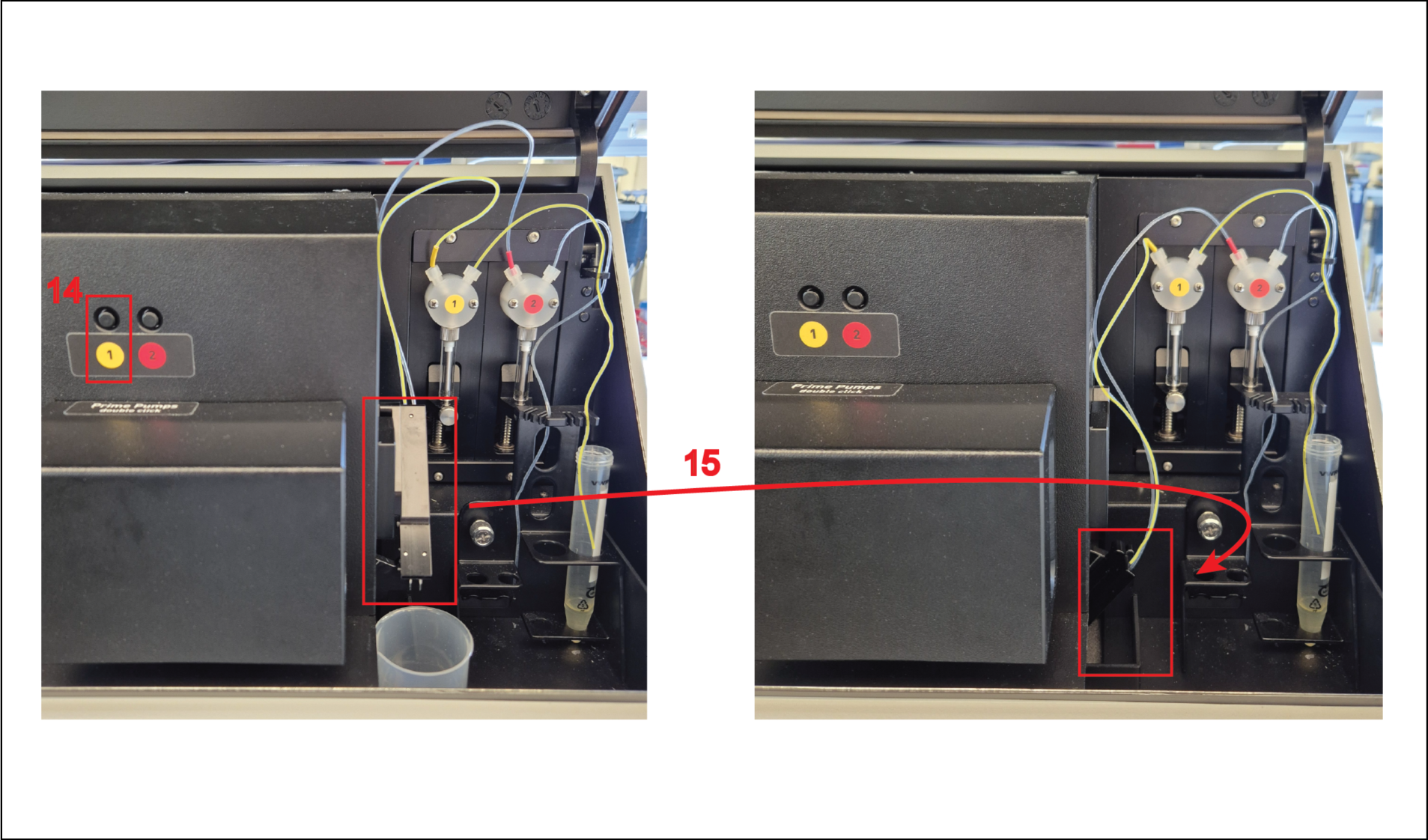

36. To measure donor- and BRET-channel, click on “**Manage Protocols**” to open the “**Test Protocols**” tab. Click on “**Luminescence**”, select the protocol “eBRET2” and click on “**Edit**”. This opens “**Luminescence - Well Mode**” window. Verify all parameters correspond to those specified in **step 27**.
37. In the “**Layout**”, select all the wells containing BRET-samples of the 96-well plate (refer to **step 31**).
38. Click on the “**Concentrations/ Volumes/ Shaking**” tab (feature 16 in **Fixed_Image_5**), type in “**10**” in the “**Start volume**” (feature 17 in **Fixed_Image_5**) for a luciferase substrate injection volume of 10 µL per well and select the pump 1 (feature 18 in **Fixed_Image_5**), which was primed for use in **step 35**. In the “**Injection Timing**” and “**Multichromatic**” tabs, keep all other parameters at default settings as specified in **step 27**. Then click “**Start measurement**” (feature 19 in **Fixed_Image_5**).
39. In the “**Start measurement**” window, input the optimal gain settings (feature 20 in **Fixed_Image_5**) and fill out the plate identification ID1, ID2 and ID3 , then click on “**Start measurement**” (feature 21 in Fixed_Image_5).

**Note:** Specify the plate identification ID1, ID2 and ID3 analogous to **step 33**.

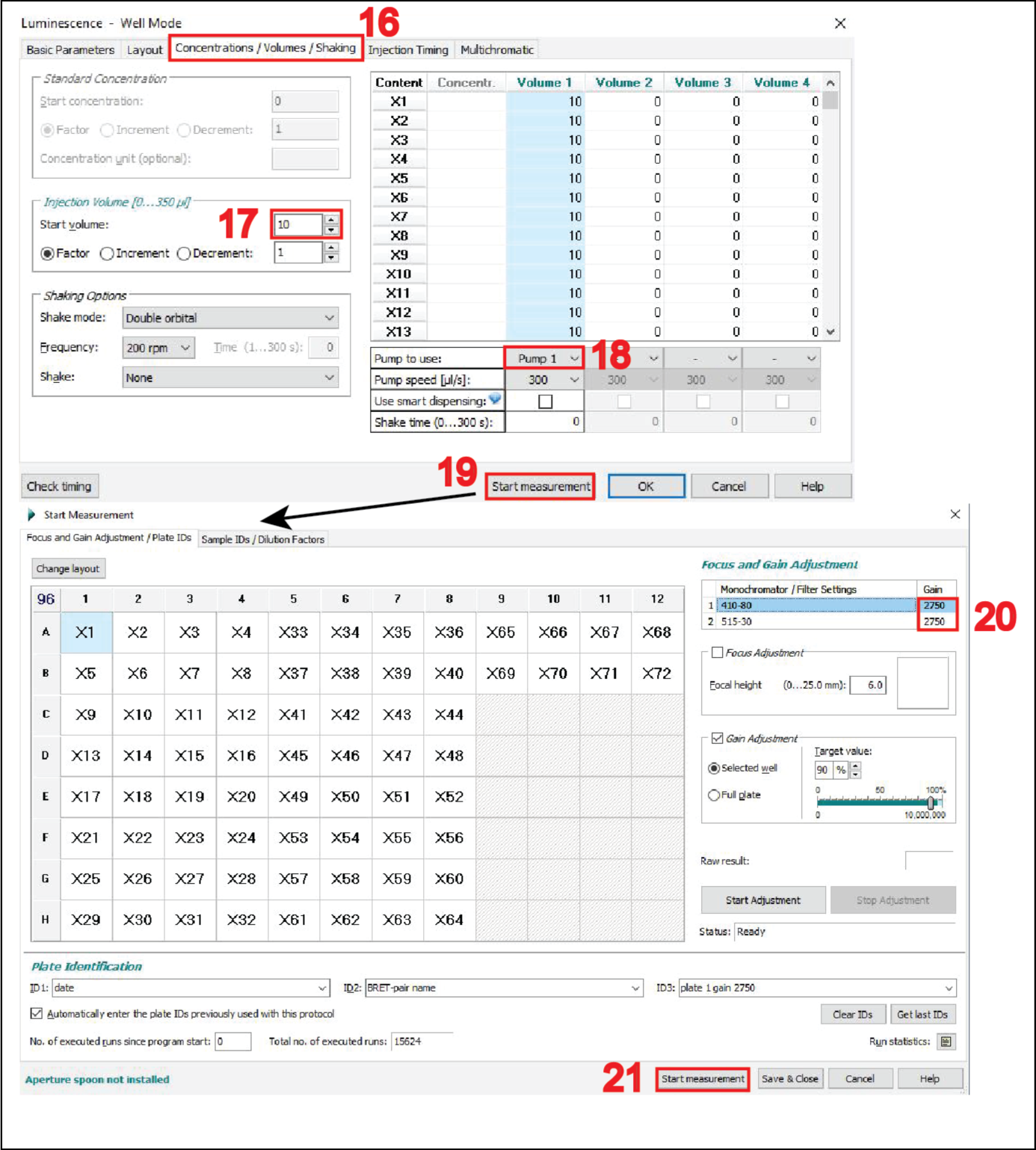

40. After the measurements, open the CLARIOstar MARS Data Analysis Software (feature 22 in **Fixed_Image_6**), where under “**Measurement Method**” either “**Luminescence**” or “**Fluorescence**” is identified. Select and open your pair of measurement files (feature 23 in **Fixed_Image_6**) then click on the small Excel icon (feature 24 in **Fixed_Image_6**) to export the displayed results as Excel workbook files.

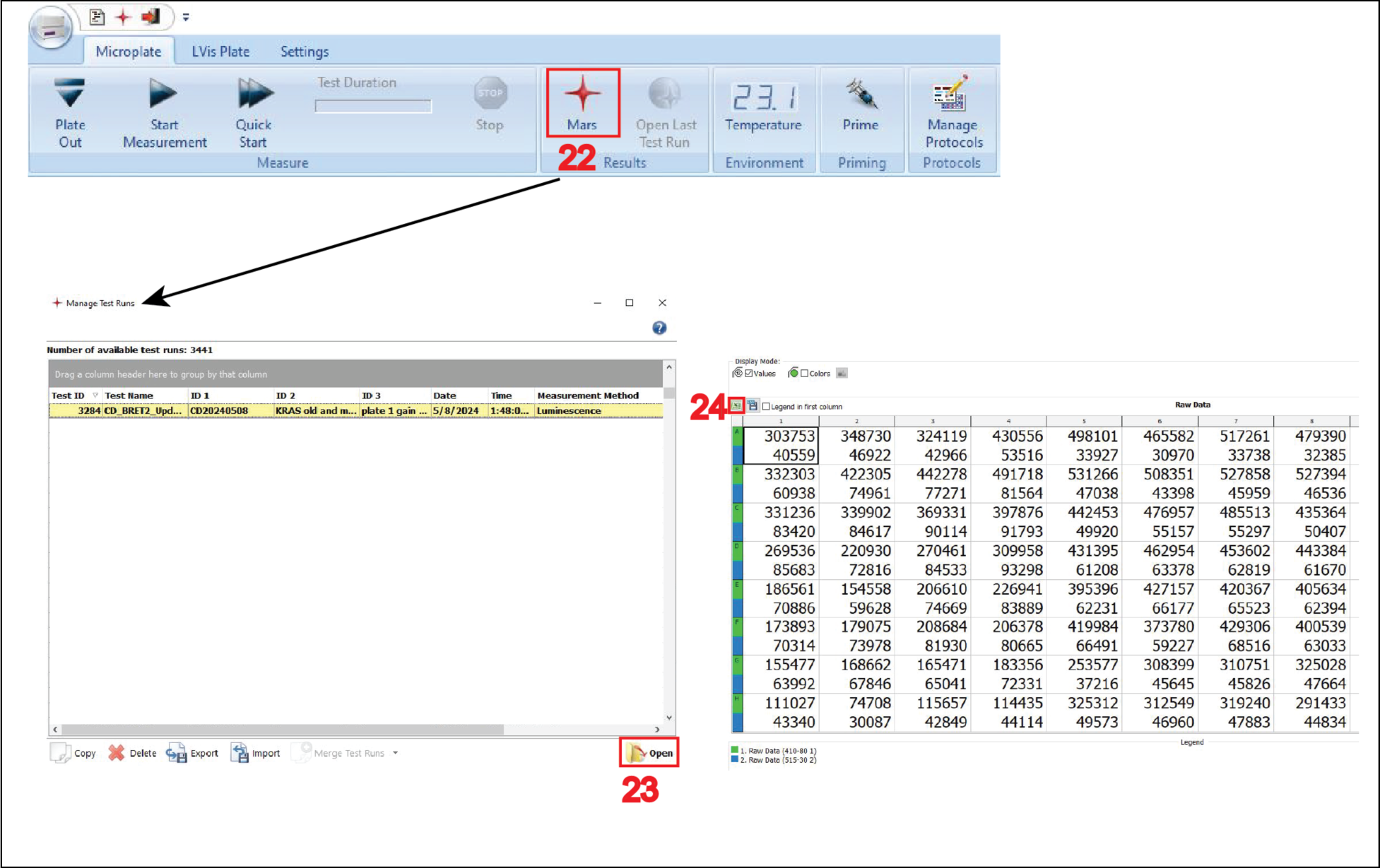

### Part 5: Data analysis in Excel and GraphPad Prism

**Timing:** 60 min

In this part, the BRET ratio, the expression signal ratio ^10^ and the normalized expression signal ratio ^11^ are calculated. Curve fitting of the data can yield the classical BRETmax and BRET50-parameters. Alternatively, we here also determine the BRETtop value, which represents the top asymptote of the BRET ratio reached within a defined acceptor/ donor range ^8^.

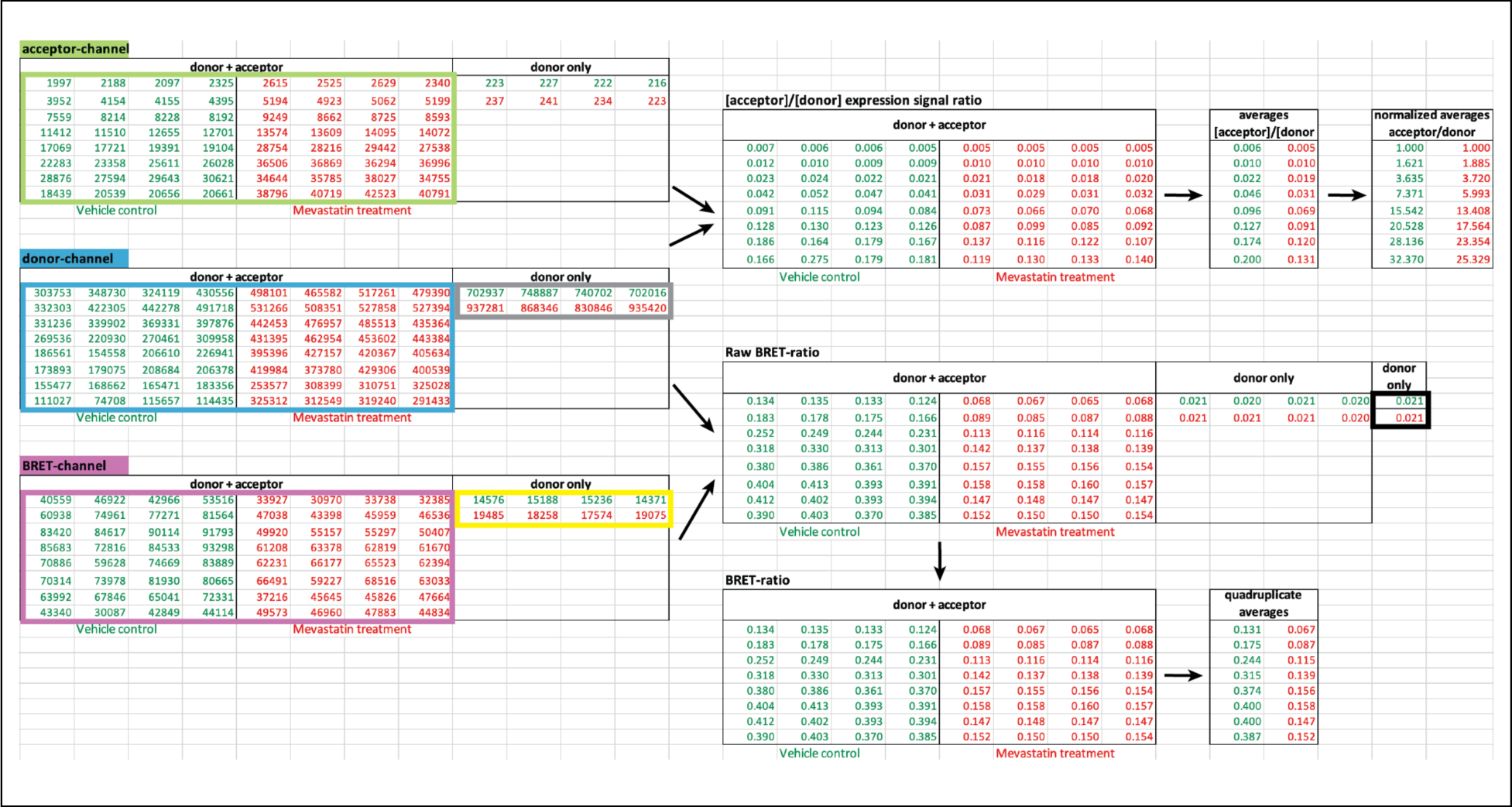

41. Merge the exported raw signals for all three channels into one Excel workbook file as shown. **Note:** The example here contains two sets of saturation-titration data as specified in **Tables 1 and S2** with vehicle control (in dark green) and mevastatin treatment (in red). Each sample is present in quadruplicates in the tables. Samples to the right, are donor-only BRET-control samples. See **<u>Troubleshooting 2</u>** for verification of raw signals in acceptor, donor-, and BRET- channels after correct gain setting.
42. The BRET ratio is calculated as the raw BRET ratio of each BRET-biosensor sample (donor + acceptor) from which the raw BRET ratio of donor-only samples is subtracted.

**Note:** The formula to obtain the BRET ratio is:

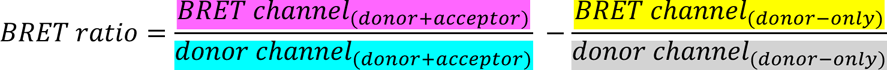

The colors in the formula refer to the colors of the boxes in the Excel sheet screenshot.

43. Therefore, calculate the raw BRET-ratio of the BRET-samples per well and subtract the average of the raw BRET ratio of donor-only samples. Then calculate from the quadruplicate technical repeats the average of the BRET ratio.
44. Next calculate the expression signal ratio per well by dividing the acceptor-channel values (boxed in light green) by the donor-channel values (boxed in blue) for all BRET-biosensor samples.
45. For the normalized expression signal ratio, further divide each expression signal ratio by the value corresponding to 1:1 A/D-plasmid ratio, here 1:1 plasmid amounts of GFP2-K-RasG12V and RLuc8-K-RasG12V.
46. Open GraphPad Prism and create an “XY” table.
47. Plot the averages across all technical and biological repeats from BRET ratio data as Y-values against the acceptor/ donor plasmid ratios from 1:1 to 40:1 (**Figure 2A**).
48. To fit the data with a hyperbolic equation, click on the “**Analyze”** tab, then under “**XY analyses**” select “**nonlinear regression**” and select “**Hyperbola** (x is a concentration)”.

**Figure 2:**
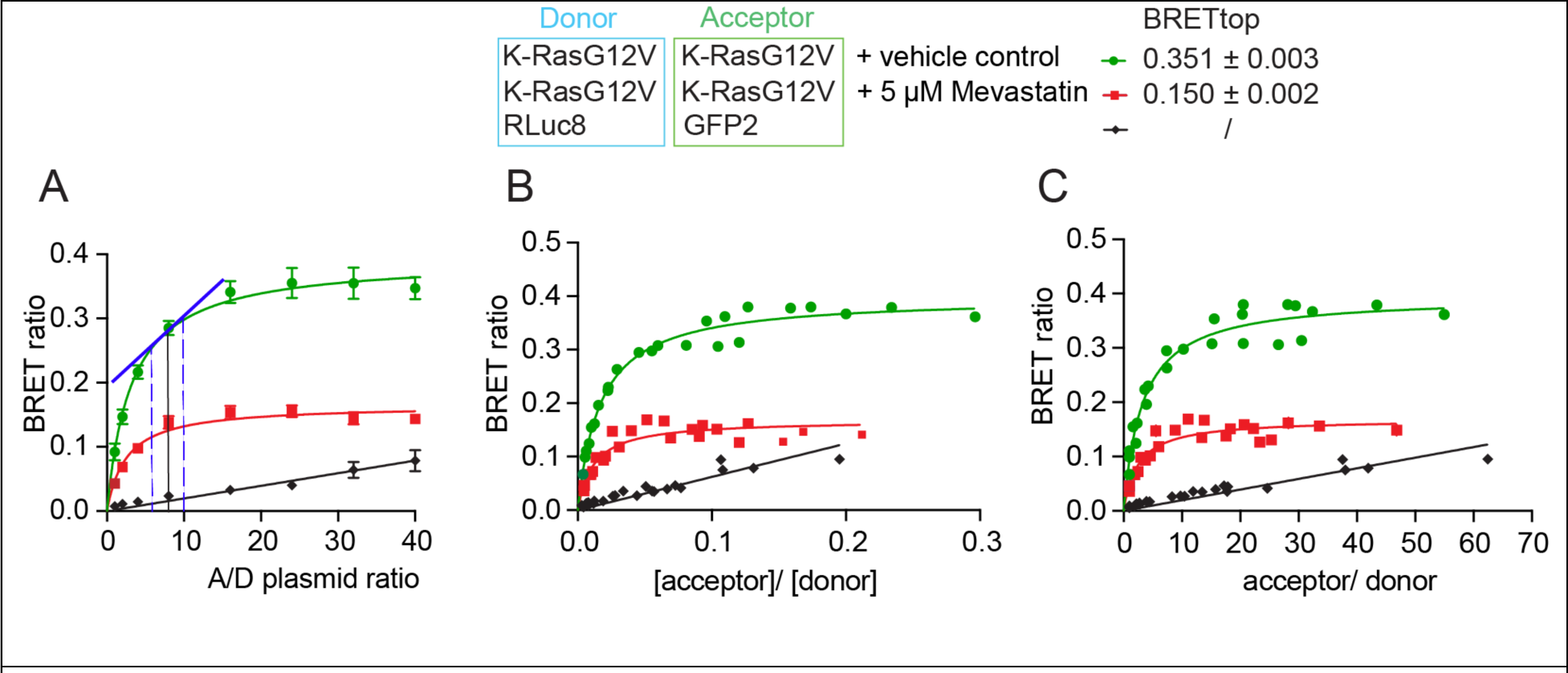
Three formats to visualize the donor saturation-titration curves of the example BRET-biosensor with and without drug treatment as compared to a background-BRET control. (**A-C**) Different representations of the same dataset from the BRET-biosensor RLuc8-K-RasG12V/ GFP2- K-RasG12V with vehicle (green), and the same with 5 μM mevastatin treatment for 24h (red). As a non- interacting control, the donor and acceptor without fused K-RasG12V were analyzed (black). Data were fit with a saturation binding curve (green, red) or linear function (black). Means ± SEM across all technical and independent biological repeats (n=3) are plotted, resulting in larger errors of BRET-ratio values (A). By contrast, in (B,C) means ± SEM of the quadruplicates from n=3 independent biological repeats are plotted individually in one plot. Note that errors are too small to be recognized (B,C). From the plot in (A) one can determine the optimal A/D-plasmid ratio for dose-response testing of e.g. a drug (not shown). The best dynamic range is found in the pseudo-linear regime of the curve, as indicated in blue (A). The preferred plot employs the ‘[acceptor]/[donor]’ expression signal ratio on the x-axis. Given that the concentration of expressed constructs is proportional to their signal, we denote their signals in squared brackets (B). From this the normalized expression signal ratio, ‘acceptor/ donor’, is derived (C). The BRETtop values were determined with a different fit function and are indicated next to the legend.

**Note:** The formula for the saturation binding curve or rectangular hyperbola is:

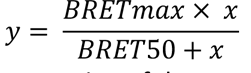

where x is a measure of the relative expression of the acceptor to the donor, and y is the BRET ratio. BRETmax represents the maximum saturation BRET signal and depends on the structural parameters (distance and orientation) of the BRET-biosensor complex. BRET50 corresponds to the acceptor/ donor ratio required to attain 50% of the maximum BRET signal and is a measure of the effective interaction probability between the interacting BRET-constructs.

49. Alternatively, use the BRET-ratio averages from technical quadruplicate repeats as y-values and plot against the averages of the corresponding expression signal ratios (**Figure 2B**) or the normalized expression signal ratios as x-values (**Figure 2C**). Fit the same hyperbolic equation (**step 48**).

**Note:** Given that the concentration of expressed constructs is proportional to their signal, we denote their signals in squared brackets, i.e. [acceptor] and [donor] for the signals acquired in the acceptor and donor channels, respectively. The expected results of raw signals from each channel are discussed in **<u>Troubleshooting 2.</u>**

50. In order to obtain the characteristic BRETtop value for data plotted as shown in **Figure 2B,C**, we employ fitting with another function. This is merely to obtain this parameter, which is not achievable with the saturation binding curve as it extrapolates the BRETmax value. First, duplicate the above data table in GraphPad Prism. In the “**Analysis**” tab, select the symbol for “**Fit a curve with a nonlinear regression**” and click on the “**one phase association**” equation.

a. In the “**Table of result**”, click on “**Nonlin fit**” in the upper left corner to open the “**Parameters: Nonlinear Regression**” tab.
b. Select “**Constrain**” and set the Y0 constant equal to 0.
c. In the “**Confidence**” section of the parameters, select “**Symmetrical (asymptotic) approximate CI**” and “**Show SE of parameters**”.
d. Go back to the “**Table of results**”. The BRETtop value is the “**Plateau**” value given with its standard error.

## Expected outcomes

To illustrate this protocol, we performed donor saturation-titration BRET experiments with the BRET-biosensor RLuc8-K-RasG12V/ GFP2-K-RasG12V (**Figure 2**). We show three plots, to illustrate the differences in appearance of the data depending on the selected x-axis. When assessing the BRET-ratio as a function of the A/D-plasmid ratio, curves appear smoothest (**Figure 2A**). This x-axis is not calculated based on actual protein expression levels but supposes that the ratio of transfected plasmid DNA is translated into corresponding protein ratios. When combining biological repeats, the uncertainty of this assumption manifests itself in a higher error of the BRET-ratio values.

From this representation one can also identify the optimal A/D-plasmid ratio (here at A/D = ∼10:1) for dose-response experiments (**Figure 2A**). Under these conditions the BRET-ratio response of the BRET-biosensor depends pseudo-linearly on manipulations that affect the interaction, such as drug-treatments ^12^.

The preferred format employs the expression signal ratio on the x-axis, as it is derived from measured signal values (**Figure 2B**). When all instrument settings are kept constant, both parameters, the BRET-ratio and the [acceptor]/[donor] expression signal ratio are values that should maintain a fixed relation between biological repeats, as they relate to actual biophysical parameters. Thus, averages of technical repeats can be visualized in one plot, without averaging biological repeat data. As an alternative in cases where the ranges of the expression signal ratio values differ much between conditions that are to be compared (e.g. different mutants of a protein that express differently), it can be advantageous to employ the normalized expression signal ratio (**Figure 2C**). However, the decision to use this representation needs to be taken in context with the specific biology.

We furthermore demonstrate that the saturation-titration curve can detect the impact of a drug treatment, here mevastatin, which prevents the lipid modification of the expressed Ras constructs and thus reduces their membrane anchorage, nanoclustering and nanoclustering-dependent BRET (**Figure 2, red curves**). With complete inhibition, all of the BRET-biosensor constructs would be cytoplasmic and should therefore behave as the tags only (**Figure 2, black curves**). Their BRET is only driven by random collisions in the cellular cytoplasm and therefore linearly depends on the acceptor/ donor ratio in the attainable expression regime. The comparison with a control, where only the tags RLuc8 and GFP2 are expressed (**Figure 2, black curves**), suggests that the mevastatin treatment does not completely inhibit membrane anchorage of all BRET-biosensors (**Figure 2, red curves**).

## Limitations

In this K-Ras-based BRET-biosensors assay, a drop in BRET such as observed by the mevastatin treatment can be due to any process upstream of Ras nanoclustering. Thus, any manipulation that impacts on Ras lipid modification, its proper trafficking or its lateral organization in nanoclusters can be detected in this assay ^13^. It is not possible to conclude that Ras or related proteins are present as dimers or other oligomers based on BRET-assay results alone.

The method can be further improved by calibrating the expression signal ratio for actual protein- stoichiometries and total expression levels. This could be achieved by using a fusion-protein of the BRET-pair with a long linker that prevents BRET as its signal ratio can be associated with a fixed 1:1 protein stoichiometry. Furthermore, using a purified acceptor protein preparation of known concentration could help to relate the signals with actual concentration equivalents.

## Troubleshooting

### Problem 1

The gain settings have not been determined accurately, eventually giving the error message “**Overflow of the signal**”, here if the signal is higher than 260,000 relative units.

### Potential solution

An optimal detector gain linearly amplifies the signal relative to the input concentration. Here we employ the transfected acceptor- (i.e. GFP2-K-RasG12V) and donor- (i.e. RLuc8-K-RasG12V) plasmid amounts as proxies for the input concentrations. Both constructs should ideally be biologically identical and correspond to the target condition to be studied i.e., here being K-RasG12V-based constructs to ensure equal expression. We individually express increasing amounts of these target BRET-biosensor constructs for the same time and using the same total DNA-amounts as later in the BRET-experiments. It is important to use actual target constructs, which will display the expression properties later found in BRET-experiments. Do not use the tags only i.e., RLuc8 and GFP2.

By following the steps described in **part 4**, we acquire a series of measurements from cells expressing the acceptor-construct using different gain settings for the acceptor channel (**Figure 3A**). The optimal acceptor-channel gain is identified as the highest gain that still linearly correlates with the amount of transfected acceptor-construct. The same will then be done for the donor channel, where the optimal donor-channel gain is identified in an analogous fashion (**Figure 3B**). For the BRET-channel we approximate the same settings as determined for the donor-channel.

**Figure 3:**
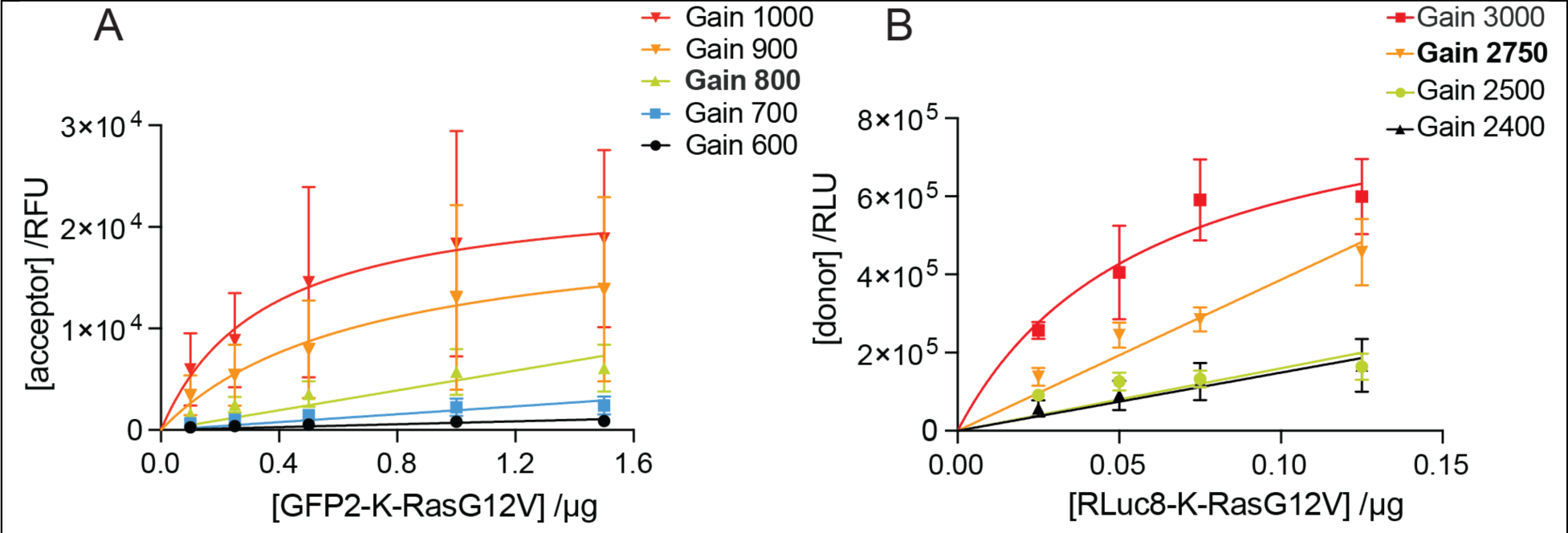
Detection channel signal gain exploration. (**A**) Acceptor-channel signals are plotted as a function of the transfected GFP2-K-RasG12V plasmid amounts at different gain values. (**B**) Donor-channel signals are plotted as a function of the transfected RLuc8-K-RasG12V plasmid amounts at different gain values. The best gain values are in bold. Curves were fit with a linear or saturation function. Plotted are means ± SEM from n = 3 independent biological repeats (A,B).

Detector gain settings usually need to be established only once for a given BRET-pair and microplate reader. Importantly, the determined gain settings have to be maintained across biological repeats that will be combined or compared. This is necessary to ascertain that the expression signal ratio scales with the stoichiometry change of expressed donor- and acceptor-constructs.

### Problem 2

Verification of raw signals in acceptor, donor-, and BRET-channels after correct gain setting (**Part 4**).

### Potential solution

The signals in acceptor, donor-, and BRET-channel should be above background and follow approximately the trends shown in **Figure 4**.

**Figure 4:**
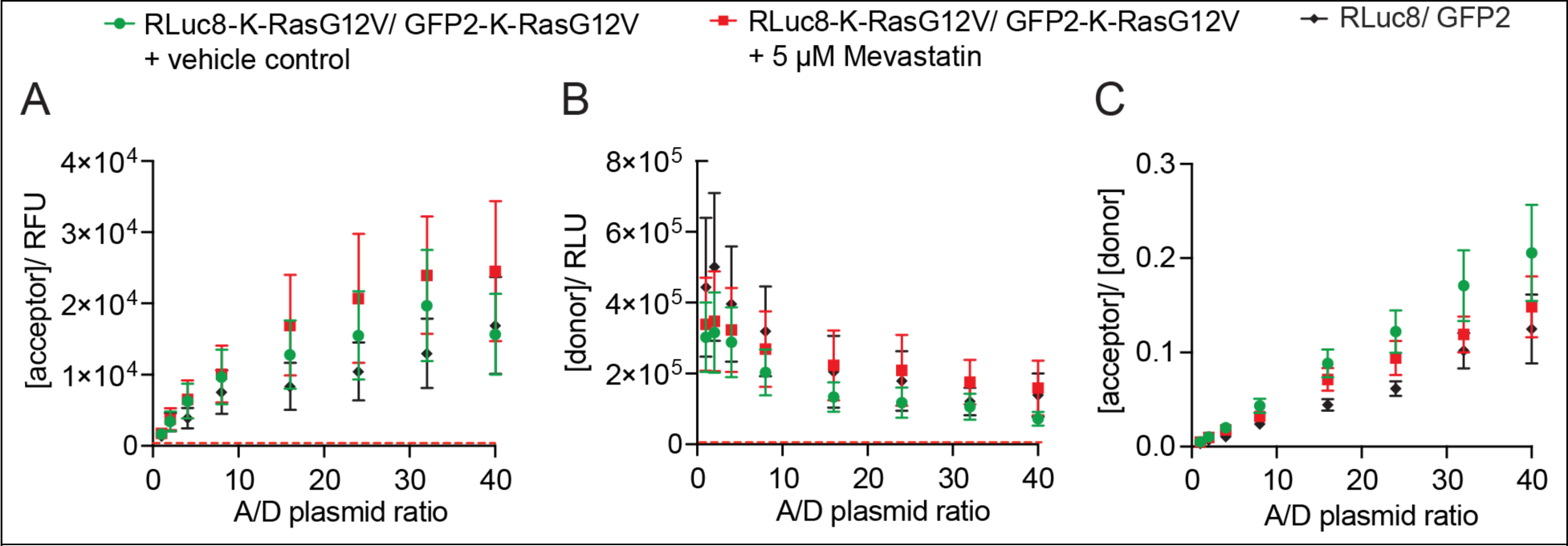
Plots to verify the good expression of the transfected BRET-biosensor constructs. (**A-C**) The raw signals from the acceptor- (A), donor- (B) and BRET-channels (C) are plotted against the A/D-plasmid ratio of transfected RLuc8-K-RasG12V/ GFP2-K-RasG12V BRET-biosensor plasmid amounts at optimal gain settings. Verify that the acceptor signal approximately linearly increases along the x-axis (A). Ideally, the signal of the donor remains constant and should be of a similar relative unit magnitude as the acceptor signals. However, here we observe that at higher A/D-plasmid ratios, the signal is reduced but somewhat constant. This is due to BRET occurring, but probably also due to the limited amount of overexpression the cell can realize. As the acceptor construct is highly overexpressed, the expression of the donor construct may somewhat be suppressed (B). The raw BRET-signal should increase with increasing A/D-plasmid ratios (C). Red dashed lines indicate the very low background signal levels of non-transfected cells. Here the acceptor-channel has merely 234 RFU and the donor-channel 34 RLU background signal. Given these values are well below those of the biosensor samples, a subtraction of this background signal from our raw signals has been omitted in our protocol.

Verify that the instrument is properly set up, notably the gain settings are correct (see **<u>Troubleshooting</u> <u>1</u>**). Start BRET-experimentation with trusted and validated BRET-biosensor constructs. The design of novel BRET-biosensors requires more experience and knowledge.

If low signals are detected, the transfection of your construct could have been insufficient, consider optimization by monitoring the expression of your BRET-constructs also by alternative means, such as flow cytometry, fluorescence microscopy or Western blotting. A low transfection efficiency can also result from sub-optimal culture conditions, such as a too dense culture or too high cell passage number. Furthermore, the cell line for heterologous expression could be relevant. It is necessary to employ a cell line with a high transfectability, as each cell should ideally be transfected with the specified A/D-plasmid ratio. We routinely use HEK293-EBNA cells for their ease of handling, transfection efficiency and high expression yields.

To increase the acceptor/ donor expression signal ratio range, consider expressing donor or acceptor construct at different ratios from those given in the example i.e., lower or higher ranges, so that the raw signals in the acceptor- and donor-channels are of similar magnitude. Alternatively, express constructs for different amounts of time e.g., express the donor-construct for a shorter time (by transfecting it later) than the acceptor-construct.

The biological impact of the BRET-construct on protein expression and cell viability or proliferation can ultimately be limiting for a successful experiment. Protein products that are toxic or cell cycle inhibitory cannot be expressed at high levels and may not be suitable for cellular BRET-measurements.

## Resource availability

- **Lead contact**: Further information and requests for resources and reagents should be directed to and will be fulfilled by the lead contact, Prof. Dr. Daniel Kwaku Abankwa.
- **Technical contact**: Questions about the technical specifics of performing the protocol should be directed to and will be answered by the technical contact, Carla Jane Duval.
- **Materials availability**: This study did not generate new unique reagents.
- **Data and code availability**: The protocol includes all datasets generated or analyzed during this study.

## Supporting information

Supplementary Information Duval et al.

## Acknowledgments

This work was supported by grants from the National Research Fund Luxembourg (FNR) INTER/UKRI/19/14174764-RAS-NANOME and INTER/NWO/19/14061736 - HRAS-PPi to DKA and AFR/17927850/Duval C./KRuptor to CJD. We are grateful to Dr. Ganesh babu Manoharan for implementing the BRET-technology in the Abankwa lab.

## Author contributions

C.L.S. participated in the initial conceptualization. C.L.S. and C.J.D. designed and carried out the experiments. C.L.S. and C.J.D. analyzed the data. C.L.S., C.J.D. and K.P. wrote the first draft of the manuscript. D.K.A. edited the manuscript and supervised the entire study.

## Declaration of interests

The authors declare no competing interests.

## Notes

### Competing Interest Statement

The authors have declared no competing interest.

