## Supplementary Information Duval et al. for "A general Bioluminescence Resonance Energy Transfer (BRET) protocol to measure and analyze protein interactions in mammalian cells"

### Supporting Information

#### Supplementary Tables

**Table S1. Example amounts of transfected siRNA a day before transfection of BRET-biosensor and donor-only (BRET-control) plasmids for donor saturation-titration BRET experiments.** "A" refers to the GFP2-tagged acceptor construct (GFP2-K-RasG12V) and "D" to the RLuc8-tagged donor construct (RLuc8-K-RasG12V). The pcDNA3.1-plasmid is used as empty vector to top up the transfected DNA amount to the same total per well.

| siRNA | well number | A/D ratio | | plasmid amounts/ ng | | | volume/ µL from 100 ng/ µL plasmid stock | | | siRNA/ nM* |
|---|---|---|---|---|---|---|---|---|---|---|
| | | A | D | A | D | empty vector | A | D | empty vector | |
| *FNTA* siRNA | 1 | 1 | 1 | 25 | 25 | 975 | 0.25 | 0.25 | 9.75 | 100 |
| | 2 | 4 | 1 | 100 | 25 | 900 | 1 | 0.25 | 9 | 100 |
| | 3 | 8 | 1 | 200 | 25 | 800 | 2 | 0.25 | 8 | 100 |
| | 4 | 12 | 1 | 300 | 25 | 700 | 3 | 0.25 | 7 | 100 |
| | 5 | 16 | 1 | 400 | 25 | 600 | 4 | 0.25 | 6 | 100 |
| | 6 | 24 | 1 | 600 | 25 | 400 | 6 | 0.25 | 4 | 100 |
| | 7 | 32 | 1 | 800 | 25 | 200 | 8 | 0.25 | 2 | 100 |
| | 8 | 40 | 1 | 1000 | 25 | 0 | 10 | 0.25 | 0 | 100 |
| | 9 BRET-control | 0 | 4 | 0 | 1 | 9 | 0 | 1 | 9 | 0 |
| negative control siRNA | 10 | 1 | 1 | 25 | 25 | 975 | 0.25 | 0.25 | 9.75 | 100 |
| | 11 | 4 | 1 | 100 | 25 | 900 | 1 | 0.25 | 9 | 100 |
| | 12 | 8 | 1 | 200 | 25 | 800 | 2 | 0.25 | 8 | 100 |
| | 13 | 12 | 1 | 300 | 25 | 700 | 3 | 0.25 | 7 | 100 |
| | 14 | 16 | 1 | 400 | 25 | 600 | 4 | 0.25 | 6 | 100 |
| | 15 | 24 | 1 | 600 | 25 | 400 | 6 | 0.25 | 4 | 100 |
| | 16 | 32 | 1 | 800 | 25 | 200 | 8 | 0.25 | 2 | 100 |
| | 17 | 40 | 1 | 1000 | 25 | 0 | 10 | 0.25 | 0 | 100 |
| | 18 BRET-control | 0 | 4 | 0 | 1 | 9 | 0 | 1 | 9 | 0 |

* The siRNA e.g. targeting the mRNA of the gene *FNTA* is transfected using Lipofectamine RNAiMAX a day before the transfection of BRET-biosensors and the donor-only BRET-control, which is not transfected with any siRNA but only receives RNAiMAX. Growth medium containing siRNA and RNA-transfection reagent needs to be removed before transfecting plasmids. Therefore, rinse cells gently once by adding 1 mL of PBS warmed to 37°C. Subsequently cells are DNA transfected as described in **part 2**. Two saturation-titration curves can then be obtained and compared, one transfected with *FNTA* siRNA and the other with negative control siRNA.

**Table S2. Example amounts of transfected BRET-biosensor and donor-only (BRET-control) plasmids for donor saturation-titration BRET experiments with drug treatment.** “A” refers to the GFP2-tagged acceptor construct (GFP2-K-RasG12V) and “D” to the RLuc8-tagged donor construct (RLuc8-K-RasG12V). The pcDNA3.1-plasmid is used as empty vector to top up the transfected DNA amount to the same total per well. Two 12-well plates are needed for the saturation-titration curves, one for the vehicle-control and one for the treatment with 5 µM mevastatin in 0.1% DMSO/ growth medium. For the mevastatin treatment, prepare a 5 mM stock solution diluted in DMSO. Take two Falcon tubes each containing 9 mL of growth medium. To the first tube you add 9 µL from the mevastatin stock. To the second tube you add 9 µL of DMSO. Vortex thoroughly to mix the solutions. Then, replace the 1 mL medium in each well with 1 mL from these treatment mixes.

| treatment | well number | A/D ratio | | volume/ µL from 100 ng/ µL plasmid stock | | | drug stock / µL** |
|---|---|---|---|---|---|---|---|
| | | A | D | A | D | empty vector | |
| vehicle control | 1 | 1 | 1 | 0.25 | 0.25 | 9.75 | 1 |
| | 2 | 4 | 1 | 1 | 0.25 | 9 | 1 |
| | 3 | 8 | 1 | 2 | 0.25 | 8 | 1 |
| | 4 | 12 | 1 | 3 | 0.25 | 7 | 1 |
| | 5 | 16 | 1 | 4 | 0.25 | 6 | 1 |
| | 6 | 24 | 1 | 6 | 0.25 | 4 | 1 |
| | 7 | 32 | 1 | 8 | 0.25 | 2 | 1 |
| | 8 | 40 | 1 | 10 | 0.25 | 0 | 1 |
| | 9 BRET-control | 0 | 4 | 0 | 1 | 9 | 0 |
| 5 µM mevastatin | 10 | 1 | 1 | 0.25 | 0.25 | 9.75 | 1 |
| | 11 | 4 | 1 | 1 | 0.25 | 9 | 1 |
| | 12 | 8 | 1 | 2 | 0.25 | 8 | 1 |
| | 13 | 12 | 1 | 3 | 0.25 | 7 | 1 |
| | 14 | 16 | 1 | 4 | 0.25 | 6 | 1 |
| | 15 | 24 | 1 | 6 | 0.25 | 4 | 1 |
| | 16 | 32 | 1 | 8 | 0.25 | 2 | 1 |
| | 17 | 40 | 1 | 10 | 0.25 | 0 | 1 |
| | 18 BRET-control | 0 | 4 | 0 | 1 | 9 | 0 |

** Drug treatment is done the day after the DNA transfection for a total of 24 hours.
